# Metabolic analysis in intact human-derived cerebral organoids by high-resolution magic-angle spinning NMR spectroscopy

**DOI:** 10.1101/2024.06.21.599412

**Authors:** Maria Alejandra Castilla Bolanos, Vorapin Chinchalongporn, Rajshree Ghosh Biswas, Colleen Bailey, Maggie Wu, Ronald Soong, Fermisk Saleh, Andre Simpson, Carol Schuurmans, Jamie Near

## Abstract

Human-derived cerebral organoids (COs) are an emerging model system for the study of human neural development and physiology. Here, we describe the assessment of metabolism in human-derived CO’s using high-resolution magic-angle spinning (HR-MAS) NMR spectroscopy. Metabolic changes during development are assessed by studying COs at various stages of maturity. Our results suggest that COs exhibit a metabolic profile similar to *in vivo* human brain metabolism, albeit with a few notable metabolic differences.

## Main

Human-derived cerebral organoids (COs) are self-organising three-dimensional cell culture systems that resemble human brain tissue and are grown in a dish. COs are an emerging and important model system for human brain research^1^. Derived from human pluripotent stem cells (hPSCs), COs take on many of the genetic and phenotypic characteristics of the individual donor, suggesting a potential role for COs in personalised medicine. For example, COs derived from patients with Alzheimer’s disease or amyotrophic lateral sclerosis exhibit pathological hallmarks of disease^2^. COs thus provide a powerful platform both to study the pathophysiology of disease, and to develop experimental treatments.

A key advantage of hPSC-derived COs is that they can be subjected to a wide range of experimental assays, including electrophysiology, histology, and ‘omics’ approaches to study the transcriptome, epigenome, or proteome, which provide a wealth of information that is typically inaccessible in human brain tissues *in vivo*. Such assays can serve as tools for studying normal cellular function and for understanding disease pathophysiology.

Assessing metabolism in COs is important to their overall physiological characterization. Metabolic assessment is commonly performed in COs using omics approaches to probe metabolic gene regulatory networks. However, these techniques are destructive and costly, and they provide only an indirect measure of metabolic status. A potential alternative method for metabolic assessment in COs is nuclear magnetic resonance (NMR) spectroscopy; a non-destructive technique that allows direct quantification of metabolites involved in different cellular and metabolic pathways. NMR offers a key advantage in that it is translational; proton (^1^H) NMR spectroscopy allows analogous measures *in vivo* and can readily be performed on most clinical MRI systems. As a result of decades of *in vivo* NMR spectroscopy research, normative concentration ranges of many key metabolites in the human brain are well established^3^. However, to our knowledge, ^1^H-NMR measures have yet to be performed in human-derived COs.

The goal of this work was to develop a high-resolution NMR approach for metabolic analyses in intact hPSC-derived COs. This approach allows metabolic assessment in COs at different stages of maturity and in CO-based disease models. To perform NMR on intact COs, we used high-resolution magic-angle spinning (HR-MAS), a technique commonly used in the analysis of intact biological tissues^4^. Briefly, for HR-MAS, the sample is placed in a cylindrical rotor, and positioned in a probe within the bore of an NMR spectrometer. The probe orients the long axis of the rotor at an angle of 54.7 degrees (the ‘magic angle’) to the main magnetic field direction (Z-axis) and spins it rapidly about this axis. Spinning the sample reduces spectral line broadening due to magnetic field inhomogeneity, dipolar coupling, and chemical shift anisotropy effects, thereby greatly improving the spectral quality and metabolic information that can be obtained from intact biological specimens^5^.

Human embryonic stem cell (hESC)-derived COs were generated using a directed differentiation protocol involving dual Smad inhibition and a spinning bioreactor^6,7^ (**Fig. 1a**). To verify the neural identity of COs prepared with this protocol, we performed immunohistochemistry, demonstrating that 124-day-old COs expressed TUJ1, a neuronal marker, and GFAP, an astrocytic marker (**Fig. 1b**). Neurons had phenotypic properties of the cerebral cortex, expressing markers of both inhibitory interneurons, such as the gamma-aminobutyric acid (GABA) A receptor alpha 1 (**Fig. 1c**), and excitatory pyramidal neurons, including vesicular glutamate transporter 1 (VGLUT1) (**Fig. 1d**). For further assessment of CO metabolic status, a bulk RNA-seq dataset collected from day 45 COs prepared with the same methodology^7^ was probed for the expression of metabolic genes of interest, including glycolytic enzymes, and enzymes involved in the synthesis and metabolism of N-acetylaspartate (NAA), hypotaurine (hTau), glutamine (Gln), and glutamate (Glu) **(Supplementary Table 1**).

**Fig. 1:**
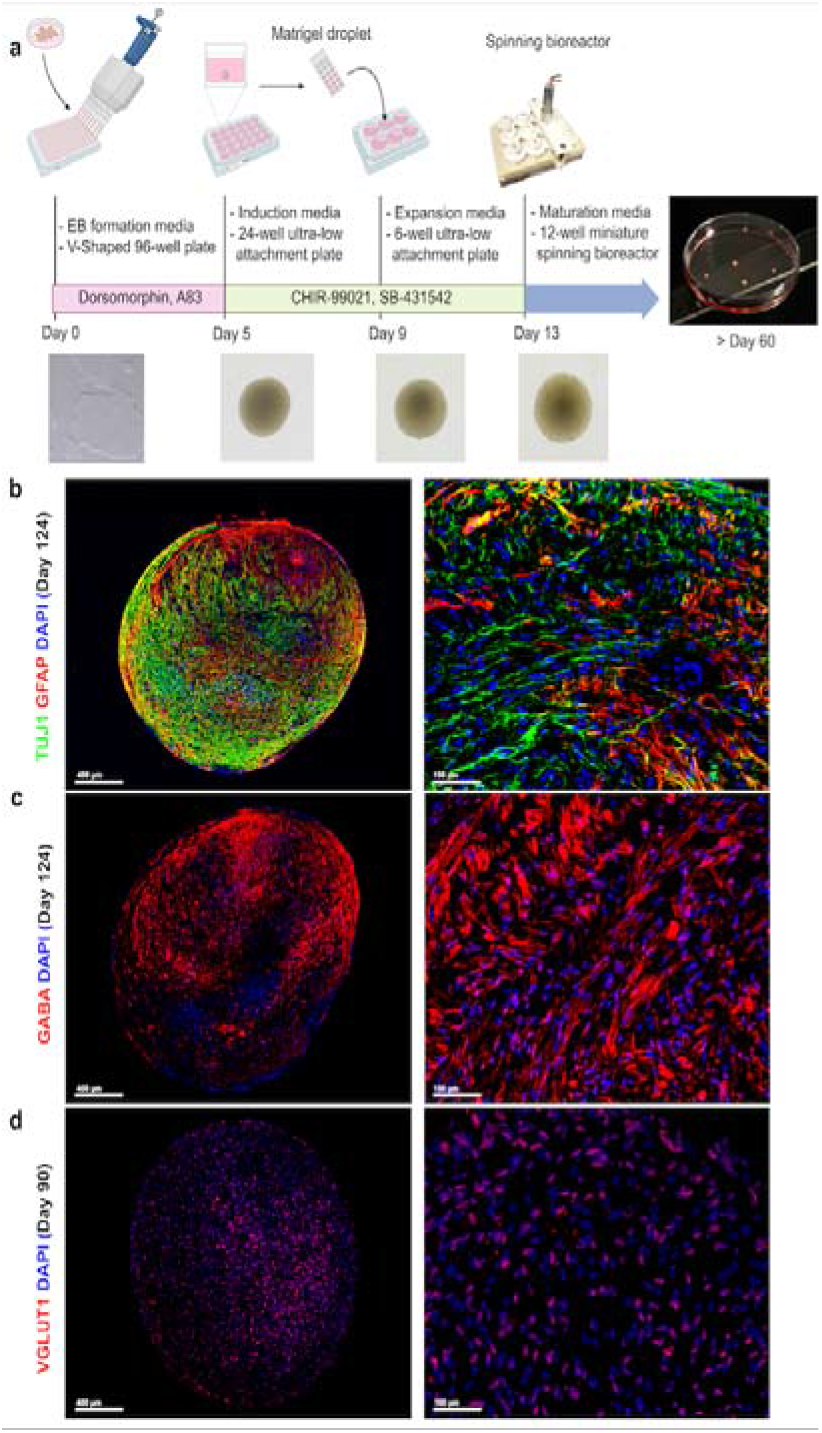
Generation and neuronal identity of hESC–derived COs. **a** Schematic overview of the generation process of cerebral organoids from hESCs using the STEMdiff^TM^ Cerebral Organoid Kit. **b, c** 124-day-old CO co-immunolabeled for TUJ1 (neuronal marker; green), GFAP (astrocytic marker, red), and GABA-A receptors (a marker of gabaergic/inhibitory neurons, red). **d** 90-day-old CO immunolabeled for vesicular glutamate transporters VGLUT1 (a marker of glutamatergic/excitatory neurons, red). Blue is the DAPI nuclear counterstain in all images. Scale bars are 400 μm (left) and 100 μm (right).

HR-MAS NMR spectroscopy was performed on a cohort of 12 identically prepared hESC-derived COs ranging from 85 to 312 days of maturity. COs (∼2 mm in diameter on average) were scanned in a 500 MHz spectrometer and maintained at 278K (5C). A spin rate of 2,500 Hz was used, as this has been shown previously to minimise the rupture of biological tissue^8^. Elements of comprehensive multiphase NMR^9^ were used to separate the NMR signals of liquid-phase small molecule metabolites from the broad spectral components of macromolecules, such as lipids and proteins. (**Supplementary Fig. 1**). In all 12 CO samples, very high NMR spectral quality was obtained (flat baselines, narrow metabolite peak linewidths of <2 Hz, **Supplementary Fig. 2**). To illustrate the CO metabolic content in aggregate, we combined the processed NMR spectra from all 12 COs into a single “average spectrum” (**Fig. 2a**). We assigned 20 individual metabolites in the upfield portion of the spectrum (0.2 - 4.6 ppm, **Fig. 2b,c**). We observed important similarities and differences between CO and *in vivo* human brain chemistry, as follows:

1. Many metabolites typically observed in the adult human brain are also present in developing COs. In most CO samples, we identified the following metabolites (in approximate order from the greatest to the smallest signal intensity): lactate, phosphocholine, glycerophosphocholine, ethanol, creatine, phosphocreatine, glycine, hTau, acetate, Gln, Glu, GABA, glucose (alpha and beta), choline, myo-inositol, alanine, valine, putrescine, lysine, and NAA. For comparison, **Fig. 2d** also shows a synthetic *in vivo* human brain spectrum which was generated using the FID-A toolkit with approximate normative human brain metabolite concentrations^3^.
2. The most abundant metabolite detected in adult human and rodent brain spectra *in vivo* is NAA, which gives rise to a prominent singlet peak at 2.0 ppm. In COs, a very low intensity NAA signal was detected in only 5 of 12 organoids (**Supplementary Figs 2-4**). This corresponded to low expression of NAA metabolic genes (NAT8L, ASPA, both with <250 normalised reads), in bulk RNAseq of day 45 COs^7^ (**Supplementary Fig. 5 and Supplementary Table 2**). NAA acts as a neuronal osmolyte, provides acetate for myelin synthesis in oligodendrocytes, and it is involved in mitochondrial metabolism^10^. The low NAA levels and gene expression observed in COs may suggest an early developmental phenotype similar to the human foetal brain, in which NAA remains low until 22-24 weeks (154-161 days) of gestational age^11^. This finding confirms that COs are potential developmental models of the human brain.
3. In the human brain, the excitatory neurotransmitter Glu is abundant, while the levels of Gln - a major metabolite of glutamate - are approximately five-fold lower (**Fig. 2b and Supplementary Figs 2-4**). In COs, the ratio of Glu/Gln appears to be reversed, or at least more balanced. In all 12 organoids, a strong glutamine resonance was observed as multiplet peaks centred at 2.41 and 2.11 ppm. In contrast, a less distinct, putative glutamate signal was observed in all 12 organoids as a multiplet at 2.32 ppm (**Fig. 2c**). Correspondingly, genes responsible for synthesis of Glu and Gln (glutaminase (GLS) and glutamine synthetase (GLUL), respectively) were expressed at similarly high levels (**Supplementary Fig. 5 and Supplementary Table 2**).
4. In 11 of 12 organoids, we detected a pair of strong triplet resonances at 2.62 ppm and 3.34 ppm, which were attributed to hTau (**Supplementary Figs 2-4**). This finding was of interest because hTau is typically not observed in human brain NMR spectra. In contrast, taurine, a closely related metabolite, is observed in both human brain spectra as well as rodent brain spectra where it is particularly abundant. We did not detect taurine in any of the CO NMR spectra. These findings corresponded to relatively high expression of CDO1 and CSAD (both related to hTau synthesis), and lower expression of FMO1, which converts hTau to Tau (**Supplementary Fig. 5** and **Supplementary Table 2**).
5. A large lactate peak was detected in most COs, in contrast to the very low lactate levels typically observed in healthy human brain tissue *in vivo* (**Supplementary Figs 2-4**). We attribute this difference to increased glycolytic metabolism in COs caused by limited oxygen supply, possibly due to poor delivery of oxygen and nutrients to the core of COs. High CO lactate levels corresponded with very high expression of a number of glycolytic genes (GAPDH, ENO1, PGK1, GPI with >3,000 normalised reads), **Supplementary Fig. 5**).

**Fig. 2:**
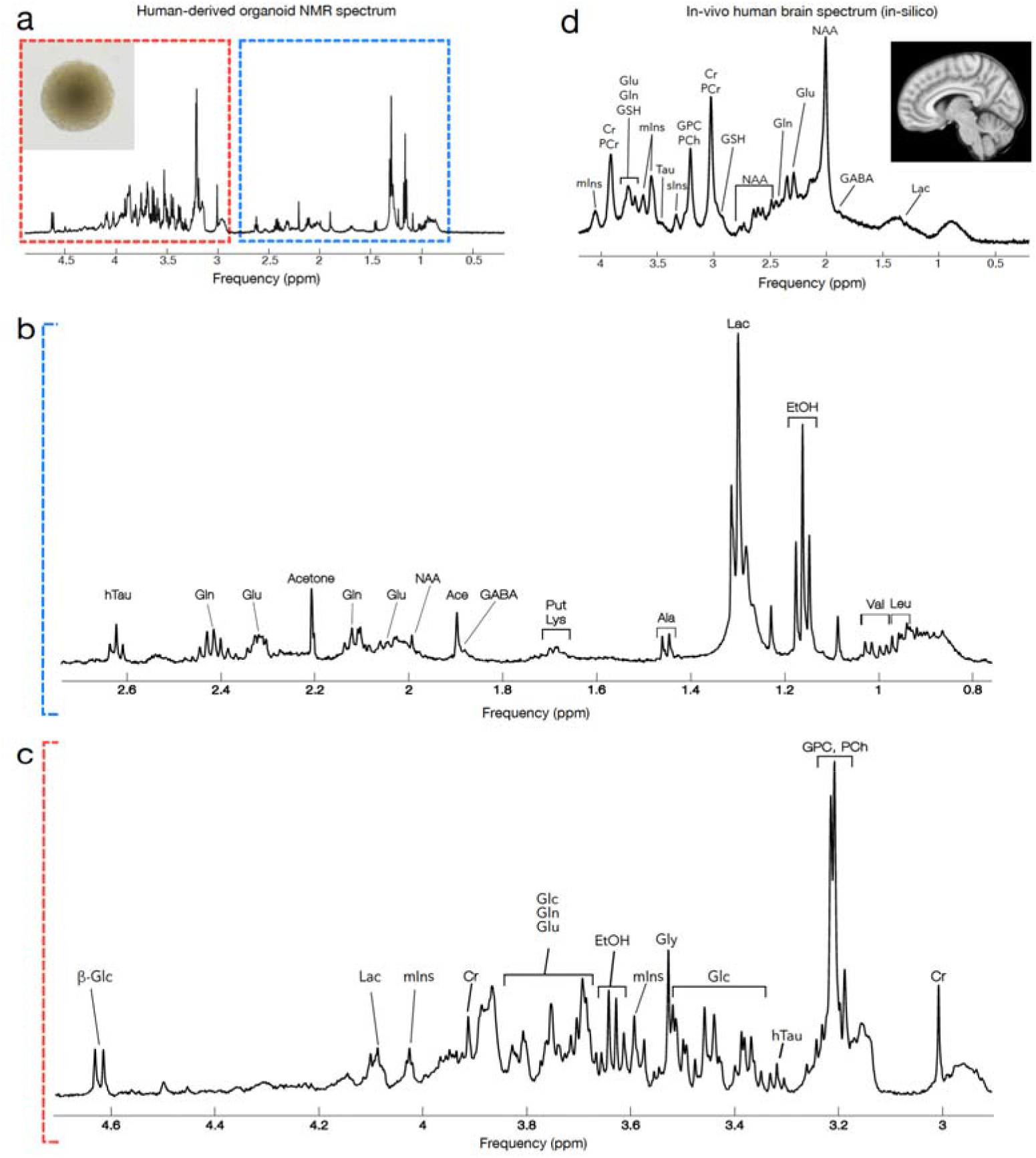
^1^H HR-MAS NMR Spectroscopy in hESC–derived COs. **a** Average spectrum of 12 hESC–derived COs from 85 to 312 days old scanned by a Bruker Avance III 500 MHz ^1^H spectrometer with a Comprehensive Multiphase NMR probe fitted with a magic angle gradient at 278 K and locked on D_2_O. **b,c** Zoomed region between 0.8–2.6 ppm and 3.0–4.6 ppm of average spectrum of 12 hESC–derived COs from 85 to 312 days of maturation. **d** *In vivo* human brain spectrum generated *in silico* indicates the most prominent metabolites in human brain tissue, including NAA, PCr, Cr, GPC, PCh, and Glu.

In summary, we developed a platform for metabolic analysis in intact hESC-derived cerebral organoids using high-resolution magic-angle spinning NMR spectroscopy. Our findings suggest that COs exhibit a metabolic profile similar to *in vivo* human brain metabolism, especially the embryonic brain, but with some notable differences in glycolysis, hTau metabolism, NAA metabolism, and Glu/Gln metabolism. We have compared our NMR metabolic measures to bulk RNA sequencing measures of gene expression for enzymes and transporters involved in these relevant metabolic pathways. This study suggests HR-MAS NMR spectroscopy as a useful methodology to assess neurochemistry and metabolism in hPSC-derived COs. Taken together, relational data analysis from these methodologies will guide efforts toward improving the engineering of human-derived COs for better modelling of *in vivo* human brains.

## Online Methods

### Cerebral organoid formation

We adapted our CO differentiation protocol from a previously published study^1^. Briefly, hESC colonies were dissociated with Gentle cell dissociation reagent (GCDR, Stem Cell Technologies, 07174) for 7 mins at 37 °C, and 12,000 cells in 100 µL STEMdiff kit EB formation media (Stem Cell Technologies, 08570) supplemented with 50 µM Rock inhibitor Y-27632 (Stem Cell Technologies, 72302) were plated in 96 well V-bottom plate (low-binding) (Greiner bio-one, 651970). This time point was called day 0 of embryoid body (EB) formation. On day 1 and 3 of EB formation, 2 µM Dorsomorphin (Stem Cell Technologies, 72102) and 2 µM A83-01 (Stem Cell Technologies, 72022) to the EB formation media. On day 5, single EBs were transfer to individual wells of 24-well ultra-low attachment plate (Corning, 3473), and media was replaced with STEMdiff kit Induction media containing 1 µM SB431542 (Stem Cell Technologies, 72234) and 1 µM CHIR99021 (Stem Cell Technologies, cat. no. 72054) and cultured for 4 more days. On day 9, COs were placed onto a single dimple of an embedding sheet (Stem Cell Technologies, cat. no. 08579) and 15 µL of Matrigel (Corning, cat. 354277) was added to each organoid to encapsulate it. Matrigel droplet-containing COs were incubated at 37 °C for 30 mins before 10-12 COs were washed into one well of a 6-well ultra-low attachment plate (Stem Cell Technologies, 38071) containing STEMdiff kit Expansion media containing 1 µM SB431542 and 1 µM CHIR99021. On day 13, individual organoids were transferred to each well of a 12-well miniature spinning bioreactor^1^ (3Dnamics (#3DNA01, Customized spin omega bioreactor-12 well) containing STEMdiff kit Maturation media. From day 30 to day 60, extracellular matrix proteins were supplemented in Maturation media by dissolving Matrigel at 1% (v/v) containing human recombinant brain-derived neurotrophic factor (BDNF; PeproTech, AF-450-02). From day 61 onwards, COs were maintained by replacing Maturation media twice a week until the experimental collection time points, as indicated.

### Human embryonic stem cells maintenance

The use of human embryonic stem cells (hESCs) was approved by the Canadian Institutes of Health Research (CIHR) Stem Cell Oversight Committee (SCOC) and the Sunnybrook Research Ethics Board (REB - PIN: 1884). hESCs (H1/WA01) were purchased from WiCell Research Institute, Wisconsin, USA. ESCs were maintained in 5% CO2 incubators at 37°C. hESCs were cultured under feeder-free conditions in mTeSR Plus media (Stem Cell Technologies, 100-0276) on plates coated with Matrigel (Corning, 354277). Versene (Thermo Fisher Scientific, 15040-066) was used to dissociate hESCs into smaller clumps by manual pipetting every 4-5 days for maintenance. hESCs were routinely tested for quality control with a qPCR-based human stem cell pluripotency detection kit (Sciencell, 0853) and hPSC genetic analysis kit (Stem Cell Technologies, 07550).

### Cerebral organoid processing and immunostaining

COs were rinsed with ice-cold phosphate-buffered saline (PBS, without Ca^2+^ and Mg^2+^) (Wisent, 311-010-CL), fixed in 4% paraformaldehyde (PFA, Electron Microscopy Sciences, 19208) in PBS at 4°C overnight. PFA was rinsed off using three washes for 5 mins in PBS before cryopreservation by immersion in 20% sucrose (Sigma, 84097)/1X PBS at 4°C overnight. COs were embedded in optical cutting temperature (O.C.T™) compound (Tissue-Tek®, Sakura Finetek U.S.A. Inc., Torrance, CA) on dry ice and stored at -80°C. 10-micron sections were collected with a Leica CM3050 cryostat (Leica Microsystems Canada Inc., Richmond Hill, ON, Canada). Samples were collected on Fisherbrand^TM^ Superfrost^TM^ Plus Microscope Slides (Thermo Fisher Scientific, 12-550-15). Cryosections of fixed COs were washed in 0.1% Triton X-100 (Sigma, T8787) in PBS (PBST), then blocked for 1 hour at room temperature in 10% horse serum (HS, Wisent, 065-150) in PBST. Primary antibodies were diluted in blocking solution as follows: TUJ1 (1:500, BioLegend #802001), GABA (1:500, Sigma #A2052), GFAP (1:500, Novus #100-53809) and VGLUT1/SCL17A7 (1:2000, Synaptic Systems #135 302). After one hour at room temperature, slides were washed 5 times for 5 mins in PBST and incubated with 1:500 dilutions of species-specific secondary antibodies (Invitrogen Molecular Probes) for 1 hour at room temperature. Slides were washed five times in PBST and counterstained with 4’,6-diamidino-2-phenylindole (DAPI, Invitrogen, D1306) and mounted in Aqua-polymount (Polysciences Inc., 18606-20). All images were taken using a Leica DMi8 Inverted Microscope (Leica Microsystems CMS, 11889113).

### NMR Spectroscopy

On the morning of the NMR experiment, the organoid to be studied was placed in a 4 mm, reduced volume (50 μL) MAS rotor (HZ07213, Bruker Ltd., Billerica, MA, USA). The remainder of the rotor volume was filled with D_2_O. Care was taken to ensure that no air bubbles were present in the rotor before putting on the cap. Rotors were then immediately transported to the NMR for scanning.

HR-MAS NMR experiments were conducted adapting protocols established by Fortier-McGill^2^. All NMR experiments were performed on a Bruker Avance III 500 MHz ^1^H spectrometer (Bruker, Billerica MA, USA), equipped with a 4 mm ^1^H-^13^C-^15^N (^2^H lock) Comprehensive Multiphase (CMP) NMR probe (Bruker Switzerland AG, Fällanden, Switzerland) fitted with a magic angle gradient. All experiments were maintained at 278 K and locked on D_2_O. Samples were spun at the magic angle at a relatively slow spin rate of 2,500 Hz to prevent the rupture of biological tissue^3^, while still positioning the spinning sidebands of water outside the spectral window.

^1^H experiments spectra were collected using 16,384 time-domain points, a spectral width of 20 ppm, 8 dummy scans and 512 total scans. Presaturation using an effective B_1_ field of 100 Hz was prepended to all experiments. Experiments used for quantitation were performed at 5 x T1 of the sample (∼11.82 seconds) between subsequent scans.

### NMR spectral editing

In NMR of biological tissue, prominent and broad resonances from macromolecules can interfere with quantification of the narrower metabolite resonances of interest. Therefore, to isolate the mobile metabolite fraction, a spectral editing approach was employed as previously described^4,5^. Briefly, the freely diffusing components (metabolites in solution) were recovered using inverse diffusion editing (IDE), whereby a strongly diffusion-weighted spectrum (DE: diffusion gradient on, depicting only macromolecular components/components with restricted diffusion) was scaled and subtracted from a non-diffusion weighted spectrum (diffusion gradient off, depicting both components with restricted and non-restricted diffusion, e.g. both macromolecules and metabolites). To minimise noise amplification due to the subtraction process, prior to scaling and subtraction we replaced the trailing noise at the end of the diffusion-weighted FID with zeros. Diffusion editing experiments were acquired using a bipolar pulse pair longitudinal encode-decode (BPPLED) sequence. Diffusion weighted acquisitions used an encode/decode gradient pulse of 2.4 ms at ∼60 Gauss/cm and a diffusion delay of 120 ms. Scaling and subtraction were performed in Matlab v.R2023b using the FID-A toolkit^6^. The resulting diffusion edited spectrum therefore depicts only the small molecule metabolite signals. An example of the diffusion editing procedure is illustrated in **Supplementary Fig. 1**.

### HR-MAS NMR metabolite identification and quantification

Metabolite assignments were performed using the Bioreference databases, version 2-0-0 to 2-0-5, AMIX (Analysis of MIXtures software package, versions 3.9.15, Bruker BioSpin, Ettlingen, Germany), and the Human Metabolome Database (HMDB, hmdb.ca^7,8^). For each metabolite to be quantified, the spectral region corresponding to an isolated (non-overlapped) resonance in the CO spectra was manually assigned as the integration region for that metabolite. The integration under the curve of each peak of interest was calculated using the TopSpin integration module^9^ (Bruker BioSpin, Karlsruhe, Germany).

The metabolite peak integrals described above were normalized against the total spectral power, obtained by integrating the full upfield spectral range, from 0.8 to 4.7 ppm. The normalized metabolite signal ratios were then corrected for the number of protons contributing to the integrated peak, as follows:

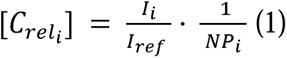

Where *C_reli_*refers to the relative metabolite concentration in arbitrary units; *I_i_* is the integral of the metabolite peak, *i,* of interest within the pre-defined integration region; *I_ref_* is the integral of the full spectral range; and *NP_i_* are the number of protons contributing to the metabolite signal of interest. **Supplementary Table 2** displays the estimated metabolite concentration in all 12 hESC-derived COs.

### Transcriptomic data analysis

Bulk RNA-seq data^10^ was mined from day 45 hESC-derived COs identically generated with the cerebral organoid formation protocol described above. For analysis of NAA, Gln/Glu, and hTau metabolic pathways, genes of interest were identified as follows. First, we searched each of the above metabolites of interest within the Recon3Map metabolic atlas, within the Virtual Metabolic Human online database (https://vmh.life) to identify key metabolic pathways involving the synthesis and degradation of each metabolite. We then noted the key enzymes and transporters (and their associated genes) involved in each of the metabolic pathways identified above. For analysis of glycolytic metabolism, a gene set was selected based on Biological Process Gene Ontology Analysis, focusing on the genes involved in the conversion of glucose to pyruvate. All genes of interest, including their metabolic roles, are listed in **Supplementary Table 1**.

To provide points of comparison for the CO gene expression findings, we studied the same genes of interest in publicly available human and rat brain transcriptomic datasets. For human transcriptomic data, we used the BrainSpan transcriptomic atlas of the developing human brain (www.brainspan.org), which provides regional RNA sequencing data from healthy human brain in multiple brain regions, and at a range of ages from 8 months post conception to 40 years of age. For our analysis, we computed an average foetal cortical expression level (computed as the average expression across all foetal timepoints from 9 to 37 weeks post conception) and an average adult cortical expression level (computed as the average expression level across all adult timepoints from 19 to 40 years of age). To obtain a single “cortical” measure, the above measures were then averaged across all cortical brain regions (dorsolateral prefrontal cortex, ventrolateral prefrontal cortex, anterior cingulate cortex, orbital frontal cortex, primary motor-sensory cortex, parietal neocortex, posterior superior temporal cortex, inferolateral temporal cortex, and occipital neocortex). For rat brain transcriptomic data, we used the Nile rat spatial transcriptomic dataset described in Toh et al^11^, which is available publicly via the gene expression omnibus (https://www.ncbi.nlm.nih.gov/geo/query/acc.cgi?acc=GSE215233). To obtain a single cortical measure, we took the average of the left caudal cortex and middle rostral cortex measures.

## Discussion

To our knowledge, this is the first study demonstrating the use of high-resolution ^1^H HR-MAS NMR spectroscopy in hPSC-derived COs. However, a few previous studies have performed other variants of NMR spectroscopy in organoids or other 3D cell models. Knitch et al^12^ used an NMR-based approach to obtain spatially resolved spectral information in 3D cell spheroids. However, unlike the current study, the cell spheroids used in the study of Knitch et al were derived from thymic NUT carcinoma cell lines, and were not neural. In agreement with our own finding of overall elevated lactate in COs, the spatially resolved approach of Knitch et al. revealed exceedingly high lactate concentrations and depleted glucose concentrations toward the oxygen-depleted centre of Ty82 cancer cell spheroids. More recently, Sapir et al. used hyperpolarized carbon-13 (^13^C) NMR spectroscopy and phosphorus-31 NMR spectroscopy to assess lactate dehydrogenase activity in COs^13^. This approach had the advantage that it enabled quantification of the metabolic rate of LDH activity. In the future, we hope to extend our approach to enable similar quantitative metabolic flux measures in hESC-derived COs using indirect ^13^C-or ^2^H-labelled glucose administration, as demonstrated previously in *in vivo* magnetic resonance spectroscopy (MRS) studies^14–17^. Lastly, van der Kemp et al.^18^ reported a comparison between HR-MAS NMR spectroscopy of intact colorectal cancer organoids, and solution NMR spectroscopy of polar extracts. While the van der Kemp et al. study finds that metabolic measures are highly correlated between intact cancer organoids and tissue extracts, the study also highlights the benefits of studying intact tissue, since it is non-destructive and enables the potential to conduct follow-up analyses such as omics analyses or histology.

NMR methods similar to those outlined here have previously been applied to the study of intact human brain tissue biopsy samples as well as other tissue specimens^19–21^. The CO spectra presented herein are similar to those presented in previous NMR studies of human biopsy tissue in terms of spectral quality. In future, comparison of NMR spectra between COs and brain tissue biopsy samples may provide further insights into key metabolic similarities and differences between organoid and *in vivo* metabolism.

A limitation of the current approach is the use of simple peak integrals for metabolite quantification. Peak integration does not allow the separate quantification of metabolites with overlapping spectral peaks, and results may be influenced by background signal contamination or spectral baseline offsets. Fortunately, the CO spectra obtained in this study had very high spectral quality with narrow linewidths (<2 Hz) and flat baselines. As a result, we were able to successfully identify and integrate at least one isolated (non-overlapping) spectral peak for each of the 16 metabolites reported. In the future, we hope to further improve the quantification of CO NMR spectra by implementing a linear combination modelling (LC model) approach as is commonly employed (and recommended^22^) in *in vivo* MRS analysis^23–27^. In contrast to peak integration, the LC modelling approach can successfully quantify partially overlapped metabolite signals, thus potentially increasing the number of metabolites that can be quantified. Although we initially attempted to use LC modelling for quantification of the CO metabolite levels in this study, this approach has so far been unsuccessful for the following reasons: In LC model analysis of *in vivo* MRS data, the individual metabolite basis functions are typically generated via quantum mechanical simulation of each metabolite’s spectral response based on prior knowledge of the chemical shifts and coupling constants of the metabolite spin system^20,28^. This prior knowledge is typically found in literature^28,29^. Due to the relatively broad linewidth (5-10 Hz) of *in vivo* NMR data, LC model analysis thereof is relatively insensitive to inaccuracies in the chemical shift and coupling constant values used for basis function simulation. In contrast, since HR-MAS NMR data have very high spectral resolution and narrow linewidths (1-2 Hz), LC modelling is highly sensitive to very small inaccuracies in the prior knowledge. Moreover, published chemical shifts and J-coupling constants are based on *in vivo* conditions (temperature, pH), which may differ from CO experimental conditions. Therefore, successful LC modelling of high-resolution NMR data will require very careful determination of the precise chemical shifts and coupling constants of each metabolite spin system of interest under the specific experimental conditions, a problem which we are currently attempting to address.

To our knowledge, this work represents the first time that relationships between human brain tissue HR-MAS NMR metabolite measures and gene expression have been described to reveal unique metabolic profiles in human COs. Four such relationships were explored:

### NAA metabolism

As described above, the finding of low NAA levels in COs was consistent with the gradual production of NAA during embryonic development. NAA serves as a neuronal osmolyte, provides acetate for myelin synthesis in oligodendrocytes, and it is involved in mitochondrial metabolism^30,31^. Low NAA levels are commonly observed during foetal brain development^32,33^, thus suggesting that COs may model an early developmental phenotype, as mentioned earlier. This finding confirms that COs are excellent developmental models and suggests that further work is required to develop organoids *in vitro* that match adult human brain metabolism, in which NAA is among the most abundant metabolites.

It is possible that late-developing cell-cell interactions between neurons, oligodendrocytes, astrocytes, and microglia are required for NAA metabolism. While neurons produce NAA, its synthesis depends on the presence of aspartate that is synthesised in astrocytes. Once produced, NAA is released for its hydrolysis to acetate and aspartate, a process mediated by aspartoacylase (ASPA) located in oligodendrocytes^34^. Oligodendrocytes slowly emerge in COs after 3 months of formation^35^. In our samples, oligodendrocytes were not identified before 124 days of maturity (assessed using immunohistochemistry with OLIG2, an oligodendroglia marker; data not shown), which might explain low expression of ASPA (54 +/- 68 normalised read) and the absence of NAA in earlier stages.

Five NAA-related genes were selected for analysis, including RIMKLB, NAT8L, SLC13A3, ASPA, and RIMKLA. As shown in **Supplementary Table 1** and **Supplementary Fig. 5,** most of the above genes are related to the breakdown or transport of NAA, while only one, NAT8L, is responsible for synthesis of NAA. In agreement with our finding of low NAA levels in CO NMR spectra, NAT8L was expressed at relatively low levels (213 +/- 64 normalised reads). The highest expressing NAA gene was RIMKLB, with 703 +/- 86 normalised reads. This gene is responsible for conversion of NAA to NAAG, suggesting that any small amounts of NAA produced are likely quickly converted by this enzyme. All of the remaining NAA related genes had an average expression of <100 normalised reads in COs, suggesting relatively low metabolic activity in this pathway in COs.

As described earlier we compared CO expression of NAA-related genes against a publicly available foetal and adult human brain spatial transcriptomic dataset (the BrainSpan Atlas of the Developing Brain, https://www.brainspan.org), and the publicly available rat brain spatial transcriptomic dataset of Toh et al.^11^. In both adult human cortex and rat cortex, where NAA levels are known to be high, NAA-related gene expression was dominated by the NAA synthesis gene NAT8L. Specifically, in adult human and rat cortex respectively, NAT8L expression was at >1.5x and >7.5x higher than any of the NAA breakdown or transport genes. This was in contrast to expression patterns in both foetal human cortex (where NAT8L expression was approximately equal to RIMKLB expression) and our own findings in COs (where NAT8L expression was three-fold lower than RIMKLB expression). This result therefore indicates that COs exhibit greater similarity to human foetal cortex rather than adult human or rat cortex.

The above comparison also shows a possible relationship between NAA levels in brain tissue and the ratio of expression of NAT8L to NAA breakdown/transport genes in COs. Future work will aim to study these relationships quantitatively.

### Glutamate/glutamine metabolism

As described above, we observed a more balanced ratio of Glu:Gln, compared with the typical ∼5:1 ratio observed *in vivo*. In the mammalian central nervous system, Glu is the primary excitatory neurotransmitter, while Gln is primarily synthesised from Glu in glial cells^36,37^. Specifically, following release of Glu into the synapse during neurotransmission, Glu is taken up by glial cells and converted to Gln, which is then recycled to neurons for subsequent re-conversion into Glu for use as a neurotransmitter.

Nine genes were selected for the analysis of Glu and Gln metabolism, including genes responsible for the conversion of Glu to Gln (GLUL), and of Gln to Glu (GLS), transport of Glu (SLC1A1, SLC1A2 and SLC1A3), the conversion of Glu to alpha-ketoglutarate (GLUD1), pyruvate (GPT), and oxaloacetate (GOT1), and Glu synthesis from glycine (GLDC). **Supplementary Table 1** and **Supplementary Fig. 5** show the expression of 9 metabolic genes involved in Glu/Gln metabolism that were targeted in this study.

Of the Glu and Glu metabolic genes selected, GLUL and GLS were most strongly expressed, with 745 +/- 107 and 783 +/- 46 normalised reads, respectively. Given that GLUL converts Glu to Gln and GLS does the opposite, the roughly equal level of expression of these two genes might suggest a reason for the unexpected balance in the observed Glu:Gln metabolite concentration ratio. If the above statement is true, then in brain tissue showing the more typical ∼5:1 Glu/Gln ratio, there would be a higher expression of GLS relative to GLUL (reflecting a preferential conversion of glutamine to glutamate). To test this hypothesis, we explored GLS/GLUL expression ratios using the same publicly available spatial transcriptomic dataset described above from BrainSpan Consortia and a rat brain dataset^11^, with the expectation that this ratio would be >>1.

Contrary to our expectations, we observed a balanced expression ratio of GLUL relative to GLS in the human brain transcriptomic data both in foetal brain (*GLS/GLUL_foetal_*: 0.65 +/- 0.44) and in adult brain (*GLS/GLUL_adult_*: 0.83 +/- 0.62) tissues, similar to the balanced expression ratio found in COs (*GLS/GLUL_COs_*: 1.05 +/- 0.43). In contrast, the rat dataset indicated a >5-fold higher expression of GLUL relative to GLS in the healthy rat brain (*GLS/GLUL_rat_*: 0.19 +/- 0.17). Therefore, the GLS/GLUL expression ratio does not appear to predict the Glu/Gln metabolite concentration ratio.

There are several factors that may result in the balanced ratio between the levels of Glu and Gln. One such factor is that astrocyte development is a later event than neuronal production^38,39^, and the astrocytes detected in the COs may be more immature than the neurons. At such an early stage of maturation of astrocytes, there may be depleted astrocyte-neuron interaction and low expression of glutaminase in neurons, resulting in a low concentration of glutamate present in COs.

Another factor is the reaction kinetics or enzyme activity levels, which do not necessarily increase in proportion to the expression of GLS and GLUL. Other potential factors to consider are the Glu/Gln interaction within COs, which can be influenced by experimental conditions, microenvironment, nutrient diffusion rates, hypoxic conditions from lack of vascularization, the presence of Gln in the cell culture medium, and other factors that extend beyond enzyme expression and functional activity. Therefore, future work should explore these interactions to fully understand the balance between Gln and Glu levels.

### Hypotaurine metabolism

As mentioned above, we observed strong triplet resonances at 2.62 ppm and 3.34 ppm, which we attributed to hTau. However, we did not detect the presence of taurine, a closely related metabolite, in any of the CO NMR spectra. The absence of taurine, and finding of high hTau levels is in direct opposition to human and rodent brain findings, in which taurine is abundant and hTau is absent. These unexpected findings suggest another major difference between CO and human brain metabolism that requires further understanding.

hTau is a derivative of cysteine, acting as a neuronal osmolyte that has antioxidant properties. In the cysteine sulfinic acid pathway, cysteine is first oxidised to cysteine sulfinic acid (also known as 3-sulfinoalanine), a process catalysed by the enzyme cysteine dioxygenase (CDO1). Cysteine sulfinic acid is then decarboxylated by cysteine sulfinic acid decarboxylase (CSAD) to form hTau^40^. The conversion of hTau to Tau is subsequently performed by the enzyme flavin-containing monooxygenase (FMO1). All three of the above genes were selected for hTau metabolic analysis, as well as the gene encoding cysteamine dioxygenase (ADO) which is responsible for the conversion of cysteamine to hTau. Our results show that COs expressed relatively high levels of both CDO1 (1911 +/- 408 normalised reads) and CSAD (591 +/- 138 normalised reads), with comparatively low expression of FMO1 (77 +/- 29 normalised reads).

**Supplementary Table 1** and **Supplementary Fig. 5** show the role and normalised reads for all targeted genes involved in hTau metabolism.

Since, in contrast to COs, both humans and rats exhibit high levels of taurine and an absence of hTau, we expected human and rat brain datatsets to show higher expression of the taurine producing gene FMO1, relative to the expression of the hTau synthesis genes CDO1, CSAD and ADO. However, contrary to our expectations, both human and rat cortex also exhibited >4-fold higher expression of each of CDO1, CSAD and ADO, relative to FMO1. Therefore, the opposing hTau/taurine ratios between COs and rat brains do not appear to be explained by the pattern of expression of the chosen genes. Thus, while our finding of low expression of FMO1 relative to genes involved in the synthesis of hTau might seem to support the high levels of hTau and the absence of taurine in CO NMR spectra, the conserved pattern in the expression of hTau genes in the human brain and rat brain with COs –where the concentrations of hTau and taurine are reversed– suggests that the chosen metabolic genes do not predict hTau and taurine levels.

One final possibility is that late-developing cell-cell interactions again contribute to the ratio of taurine and hTau. The taurine synthetic pathway is proposed to be incomplete in astrocytes and neurons, and metabolic cooperation between these cell types is reportedly needed to complete the pathway^41^. Accordingly, it is reported that the co-culture between neurons and astrocytes can 10-fold decrease the hTau:taurine ratio with the presence of astrocytes increasing neuronal taurine content. While we reported the presence of spontaneous development of astrocytes in COs, it is possible that poor colocalization of astrocytes and neurons (**Fig. 1**) favours the production of hTau over taurine^42^.

### Glycolysis

The finding of high NMR lactate levels in COs points towards upregulation of glycolysis. Glycolysis is the main pathway to produce energy from glucose, resulting in pyruvate and lactate as end products. The glycolytic pathway consists of more than 10 metabolic conversion steps, each with an associated enzyme/gene. A total of 15 glycolytic genes were selected for analysis (**Supplementary Fig. 5**.), and the normalised expression of these metabolic genes is available in **Supplementary Table 1.** Among these are the hexokinases (HK1 and HK2), which catalyse the phosphorylation of glucose to form glucose-6-phosphate (G6P), and the lactate dehydrogenases (LDH1 and LDH2), which catalyse the reaction between pyruvate and lactate^43^. All 15 glycolytic genes were expressed in COs, the lowest being HK1 (557 +/- 126 normalised reads), PKM (17+/- 18 normalised reads), and PKLR (4 +/- 2.57 normalised reads). PKM and PKLR encode pyruvate kinase, an enzyme responsible for ATP and pyruvate production. Interestingly, their expression levels are lower than the other glycolytic genes. Considering that pyruvate in fast exchange with lactate such that the levels of these two metabolites are generally believe to be tightly coupled, the low expression of PKM and PKLR contrasts with the high level of lactate in all 12 CO NMR spectra (**Supplementary Figs 2-4**).

The most highly expressed gene was glyceraldehyde-3-phosphate dehydrogenase (GAPDH) with a very high expression of 22220 +/- 3287 normalised reads, followed by enolase-1 (ENO1), with a high expression of 14818 +/- 4362 normalised reads and phosphoglycerate kinase (PGK1) with a high expression of 4710 +/- 1371. GAPDH is responsible for an intermediate step of glycolysis in which glyceraldehyde-3-phosphate is converted to 1-3-bisphosphoglycerate. ENO1 is responsible for a later step in which 2-phosphoglycerate is converted to phosphoenolpyruvate. In addition, PGK1 is responsible for the conversion of 1-3-bisphosphoglycerate to 3- phosphoglycerate, immediately after GAPDH. It is not clear why GAPDH and ENO1 had such particularly high expression relative to other glycolytic genes, but the overall high expression of genes in the glycolytic pathway expression remains a key finding.

Some genes in the glycolytic pathway are commonly associated with foetal brain metabolism^44,45^. Indeed, upregulation of glycolysis is often a feature of rapidly growing tissues, as they tend to quickly outgrow their oxygen supply. However, analysis of the spatial transcriptomic datasets from the BrainSpan Consortia indicated that there are higher expressions of glycolytic genes (GAPDH, ENO1, PGK1, LDHB, GPI, LDHA, ENO2, PGAM1, ALDOC, HK1, and TPI1) in adult brains compared to foetal brains. Only three of 15 genes, which are PFKL, involved in the Conversion of D-fructose 6-phosphate to fructose 1,6-bisphosphate; HK2, responsible for the phosphorylation of glucose to glucose-6-phosphate; and PFKFB3 that synthesises fructose-2,6-bisphosphate, remained at a higher expression in foetal brains.

While the distinct pattern of the high expression of GAPDH and ENO1 remains similar in human brain and COs, the rat brain does not conserve high expression of ENO1 in their glycolytic profile, which might suggest COs as a potential platform to study glycolysis in the human brain.

Genes involved in the glycolytic pathway are also strongly associated with cellular stress and therefore studied in carcinoma studies^46^. In the context of the current study, the high expression of GAPDH, ENO1, PGK1 might be due to the lack of vascularization of COs, resulting in hypoxia in the inner core of COs. Therefore, hypoxic conditions might increase the activation of genes associated with cellular stress and glycolysis.

### Ethanol NMR signal

In all CO spectra, we observed a large ethanol peak around 3.5 ppm. This was unexpected, as ethanol is normally not detected in *in vivo* MRS of human and animal brain (except following alcohol consumption). While the reason for this ethanol peak is uncertain, it may be related to the presence of an alcohol analog, 2-mercaptoethanol, in the organoid generation kit used.

## Conclusion

In summary, HR-MAS NMR performed on intact hESC-derived COs yields detailed spectra with new insights into the metabolic status of COs. Important similarities were observed between COs and the known neurochemistry of human brain tissue, highlighting the potential of this technique. As new CO protocols and disease models come online, HR-MAS NMR can identify disease-modifying metabolic differences. A more detailed omics analysis may also identify how the activity of specific metabolic and enzymatic pathways is related to observed metabolite levels. Such relational data analysis from these methodologies will direct the design of enhanced human-derived COs as models human brain metabolism.

## Acknowledgements

This work is supported by the Canadian Institutes for Health Research (JN, AS and CS, Grant #: PJT-183715).

## Supplementary Information

**Supplementary Fig. 1:**
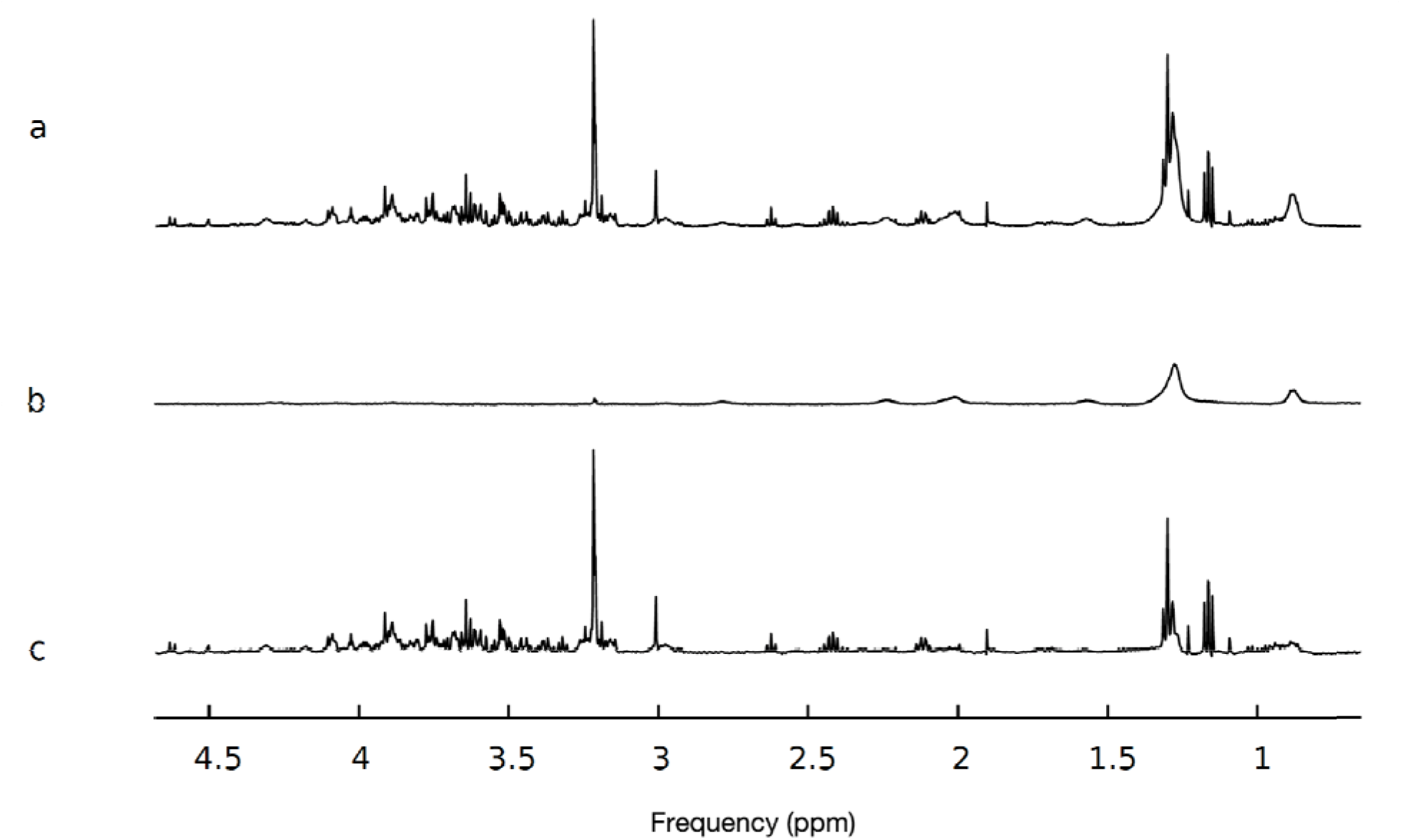
Diffusion editing for removal of macromolecular resonances in cerebral organoids spectra. **a** Original, non-diffusion-weighted spectrum showing both metabolite and macromolecular peaks. **b** Diffusion-weighted spectrum showing only macromolecular components/components with restricted diffusion. **c** Inverse Diffusion-edited spectrum obtained by the subtraction of spectrum b from spectrum a. The Inverse Diffusion-edited spectrum consists of only freely-diffusing metabolites.

**Supplementary Fig. 2:**
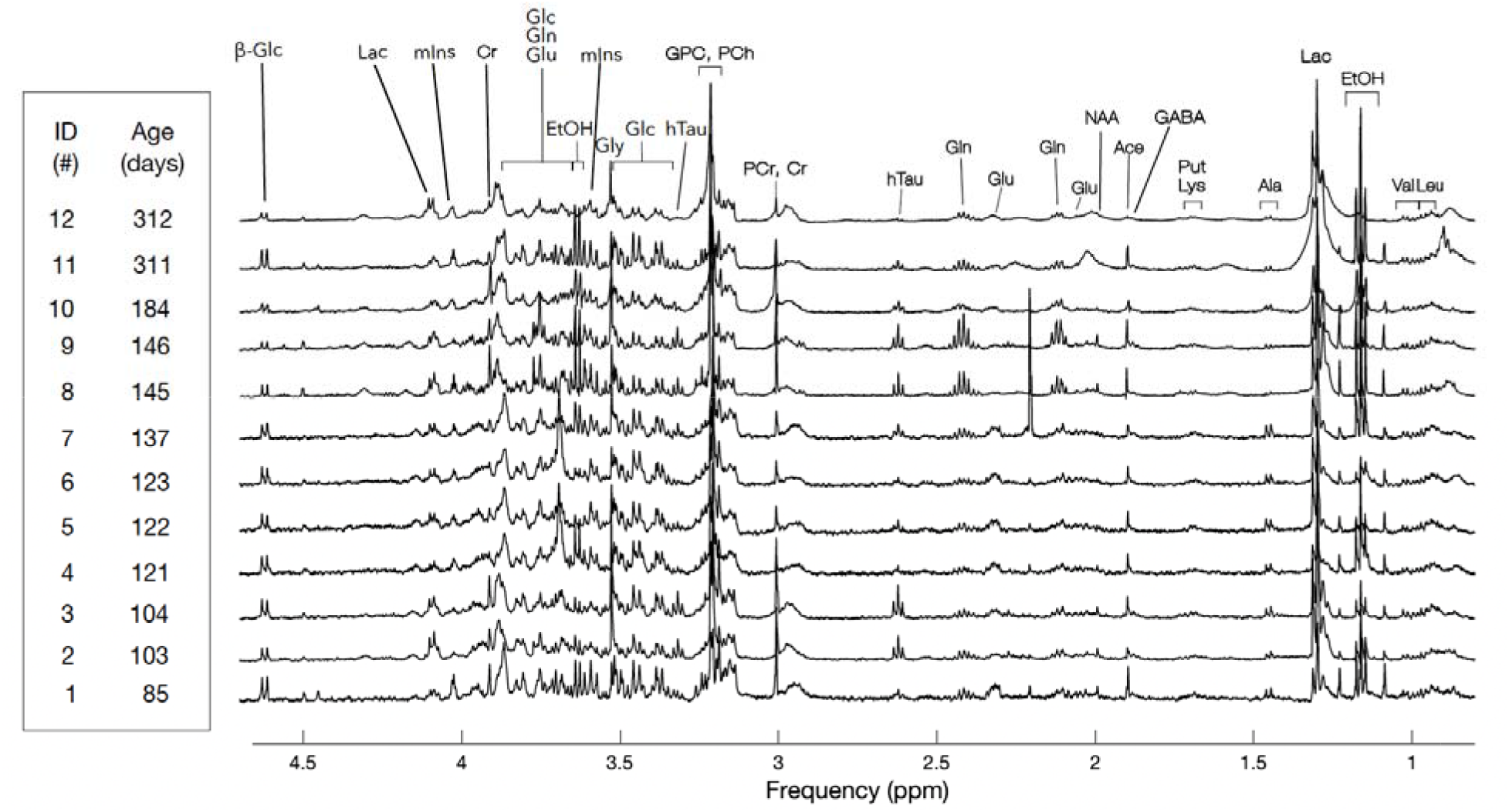
^1^H HR-MAS NMR Spectroscopy in hESC–derived COs. Spectra of 12 intact hESC–derived COs aged between 85 and 312 days. Spectra were acquired using a Bruker Avance III 500 MHz ^1^H spectrometer with a Comprehensive Multiphase NMR probe fitted with a magic angle gradient. Samples were maintained at 278 K for the duration of the experiment and were spun at 2,500 Hz. Labelling above the spectra indicates the metabolite peak assignments. Each spectrum was aligned and scaled to have the same creatine signal intensity. β-Glc: glucose; Lac: lactate; mIns: myo-inositol; Cr: creatine; Gln: glutamine; Glu: glutamate; etOH: ethanol; Gly: glycine; hTau: hypotaurine; GPC: glycerophosphocholine; PCh: phosphocholine; PCr: phosphocreatine; NAA: N-acetyl-aspartate; Pt: putrescine; Lys: lysine; Ace: acetate; GABA: γ- hydroxybutyric acid; Ala: alanine; Val: valine; Leu: leucine.

**Supplementary Fig. 3:**
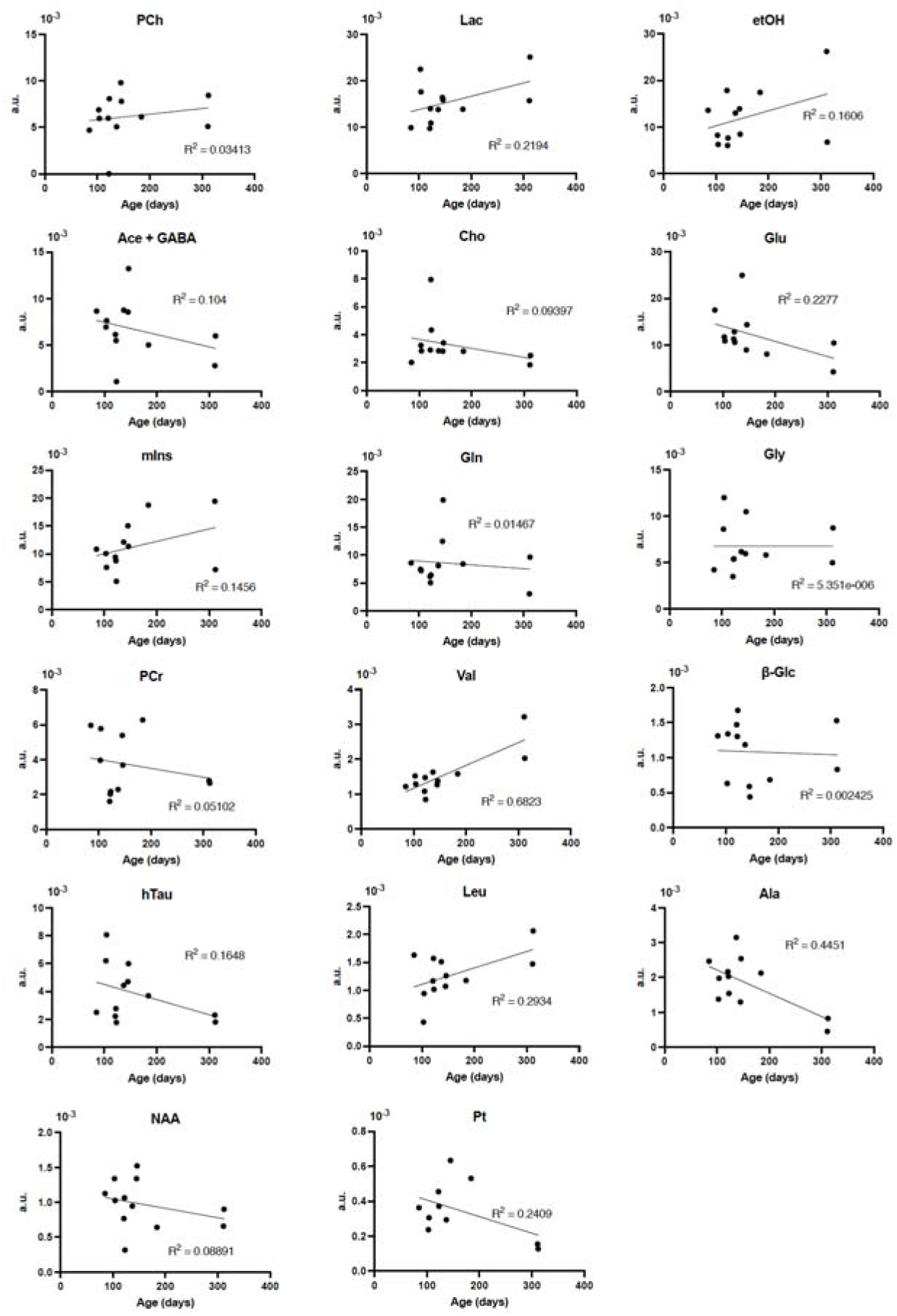
Progression of metabolite concentrations during hESC-derived CO development. Concentration of metabolites over time was recorded in 12 intact hESC-derived COs scanned between 85 and 312 days post generation. PCh: phosphocholine; Lac: lactate; etOH: ethanol; Ace + GABA: acetate and GABA; Cho: choline; Glu: glutamate; mIns: myo-inositol; Gln: glutamine; Gly: glycine; PCr: phosphocreatine; Val: valine; β-Glc: glucose; hTau: hypotaurine; Leu: leucine; Ala: alanine; NAA: N-acetyl-aspartate; Pt: putrescine.

**Supplementary Fig. 4:**
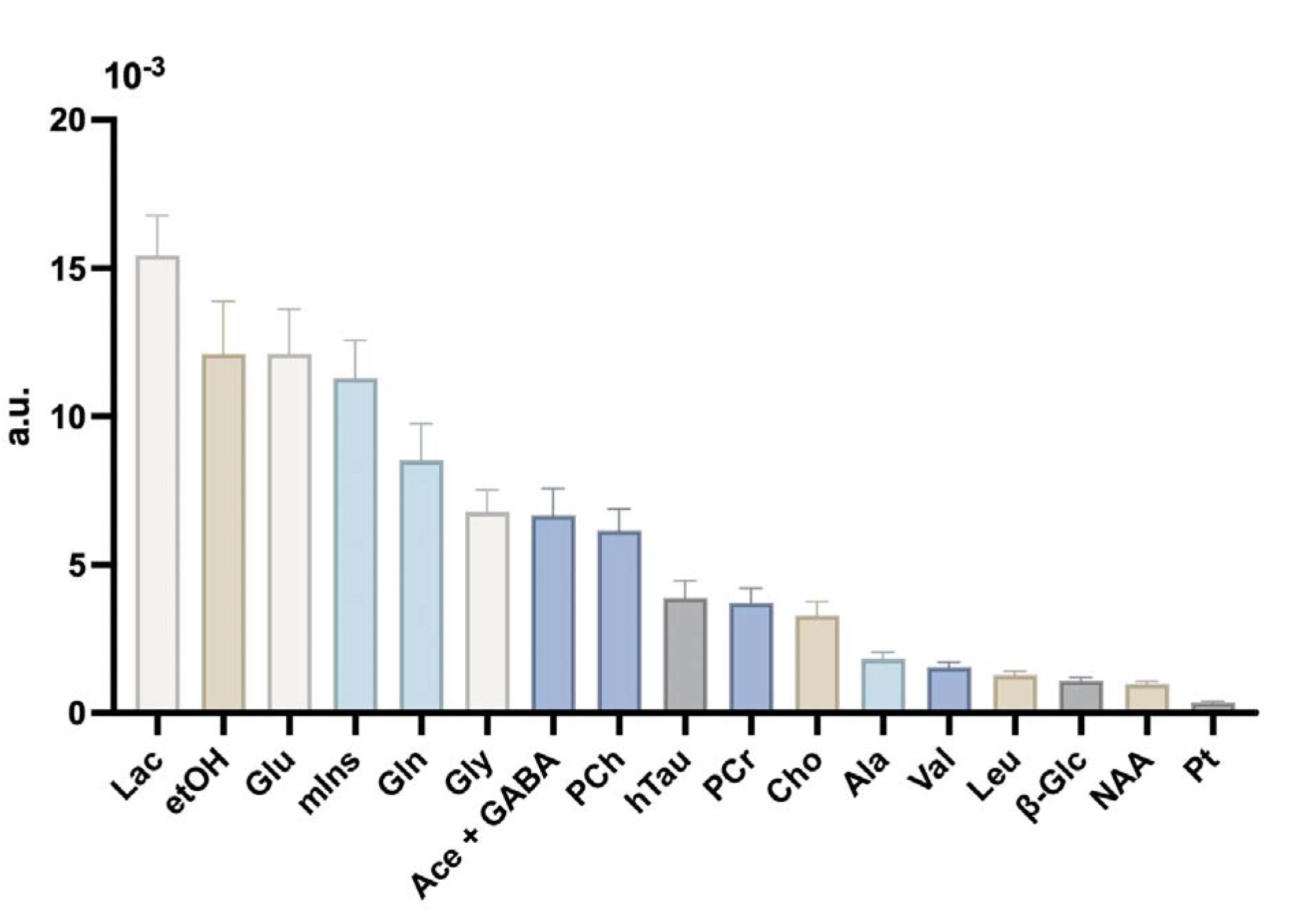
Average metabolite concentrations in hESC-derived COs. Average concentrations of all detected metabolites were recorded in 12 intact hESC-derived COs scanned between 85 and 312 days post generation. Data represent the means ± SEM for 12 independent experiments. Lac: lactate; etOH: ethanol; Glu: glutamate; mIns: myo-inositol; Gln: glutamine; Gly: glycine; Ace + GABA: acetate and GABA; PCh: phosphocholine; hTau: hypotaurine; PCr: phosphocreatine; Cho: choline; Ala: alanine; Val: valine; Leu: leucine; β-Glc: glucose; NAA: N-acetyl-aspartate; Pt: putrescine.

**Supplementary Fig. 5:**
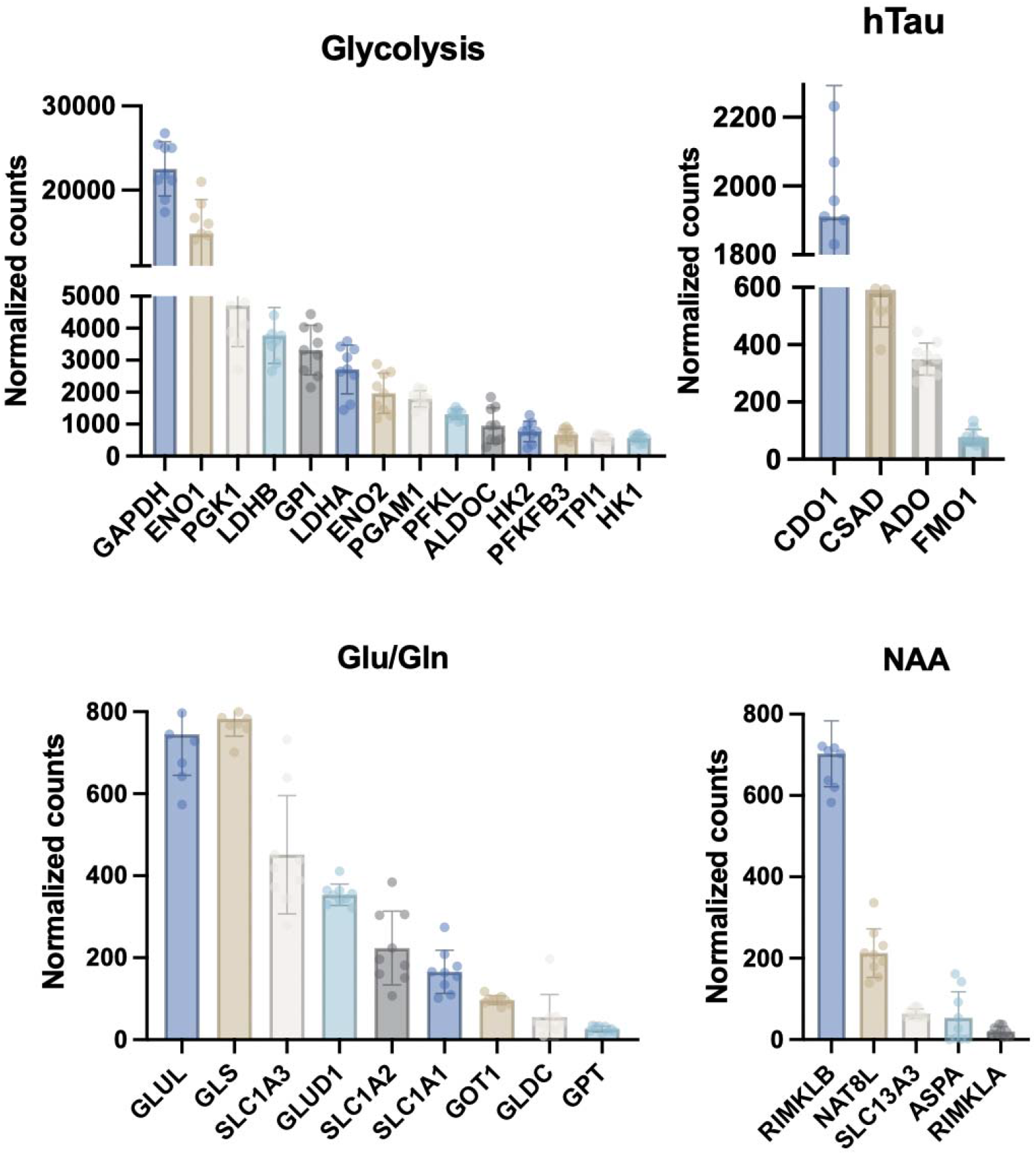
Gene expression in hESC-derived COs. Average expression of targeted genes involved in glycolysis, hTau metabolism, Glu/Gln metabolism, and NAA metabolism were recorded from 45-47 day old intact hESC-derived COs using bulk RNA-sequencing. Data represent the normalised count means per gene ± SD for independent CO experiments (n=8).

**Supplementary Table 1:**
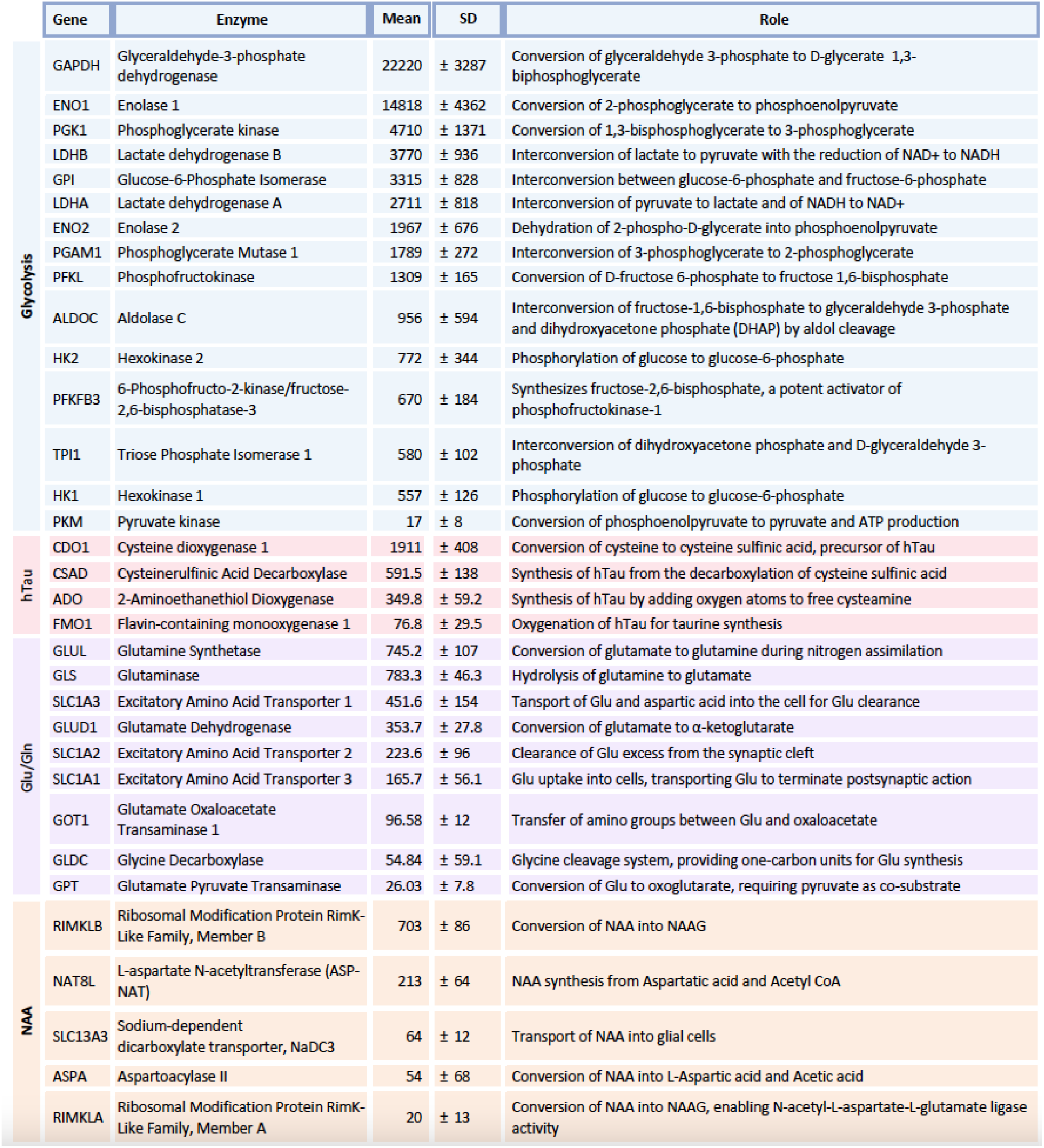
average gene expression in hESC-derived COs.

**Supplementary Table 2:**
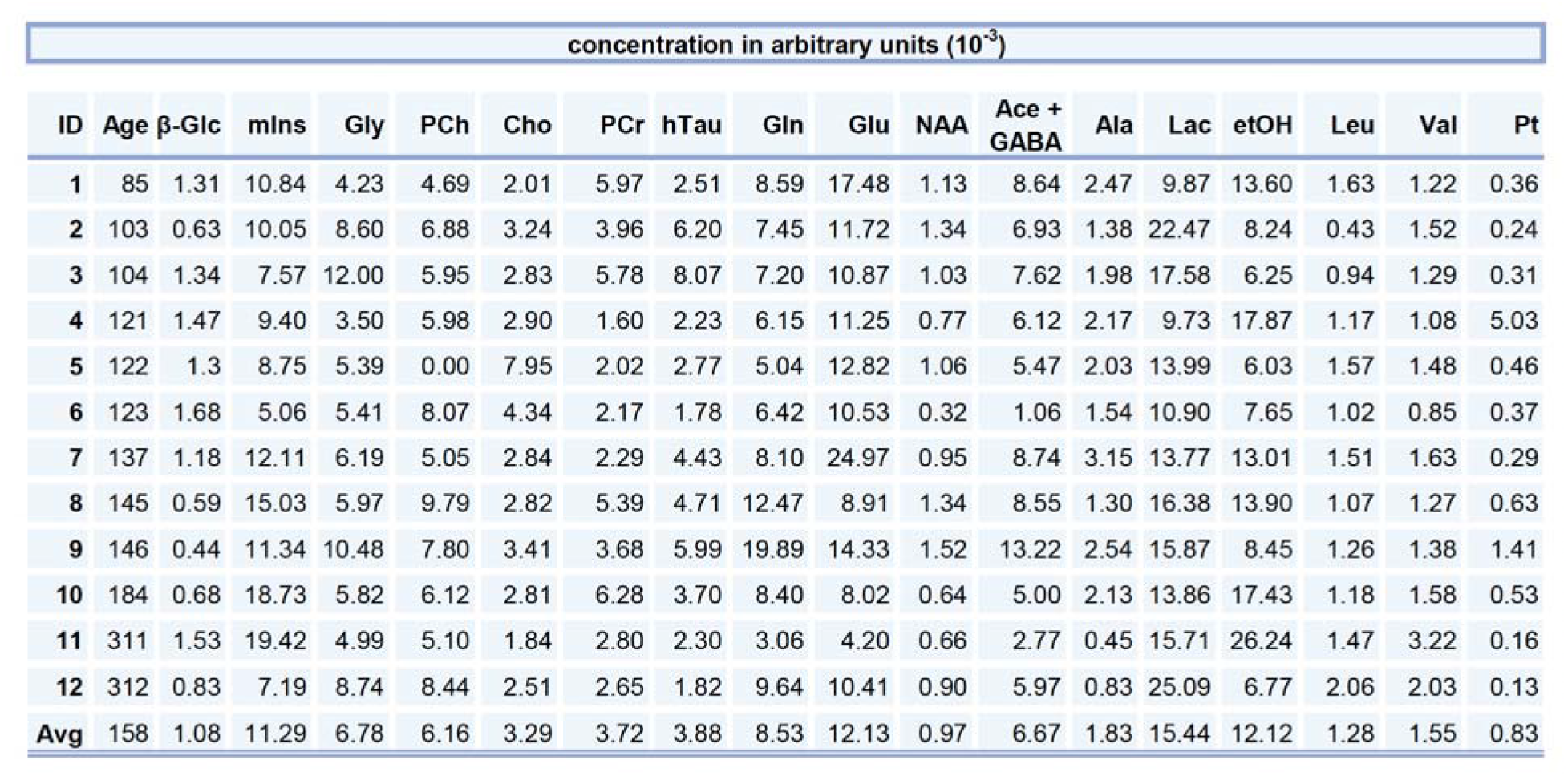
quantification of metabolites in COs at different stages of maturation.

## Notes

### Competing Interest Statement

The authors have declared no competing interest.

